# Overhauling a faulty control in the CDC-recommended SARS-CoV-2 RT-PCR test panel

**DOI:** 10.1101/2020.06.12.147819

**Authors:** Rob J. Dekker, Wim A. Ensink, Selina van Leeuwen, Han Rauwerda, Timo M. Breit

## Abstract

To battle the COVID-19 pandemic, widespread testing for the presence of the SARS-CoV-2 virus is worldwide being employed by specific real-time RT-PCR (rRT-PCR) of viral RNA. The CDC has issued a recommended panel of PCR-based test sets that entail several primer/probe sets that target the SARS-CoV-2 N-gene, but also one that targets the human RNase P gene (h-RP) as a positive control for RNA extraction and/or reverse-transcription (RT) efficacy.

We discovered that the CDC-recommended h-RP primer/probe set has a faulty design, because both PCR primers are located in the same exon, which allows for unwanted PCR-amplification of background genomic DNA (gDNA). By removing RNA from nose-swab samples by an RNase treatment, we showed that the presence of gDNA in samples resulted in false-positive signals for the h-RP test control. This is rather serious, because it could lead to false-negative test outcomes, since the CDC interpretation of an absent SARS-CoV-2 rRT-PCR signal plus a positive h-RP rRT-PCR signal is interpreted as “2019-nCoV not detected”, whereas a false-positive h-RP rRT-PCR signal resulting from amplification of gDNA should be interpreted as “Invalid Result” and the procedure should be repeated.

In order to overhaul the faulty h-RP rRT-PCR primer/probe set with minimal modification, we designed and tested several new h-RP reverse primers. Replacement of the CDC-recommended PCR reverse primer with our selected exon-exon junction reverse primer corrected the problem of false-positive results with this important SARS-CoV-2 RT-PCR test control and thus eliminated the problem of potential false-negative COVID-19 diagnoses.

## INTRODUCTION

Managing the response to the COVID-19 pandemic caused by the severe acute respiratory syndrome coronavirus 2 (SARS-CoV-2) relies heavily on massive testing of (pre)symptomatic individuals^1,2^. Currently, this is normally done by showing the presence of SARS-CoV-2 RNA in a frontal nose swab using a real-time RT-PCR (rRT-PCR) test^3^. Generally, these SARS-CoV-2 rRT-PCR test panels have several targets in one or more genes of the virus, complemented with several positive controls and negative controls for checking the correct execution of the whole procedure, including RNA extraction and RNA reverse-transcription efficacy (RT)^4^. Early 2020, the Division of Viral Disease of the Centers for Disease Control and Prevention (CDC, Atlanta, USA) has put a rRT-PCR diagnostic panel together for the detection of the 2019-Novel Coronavirus (https://www.fda.gov/media/134922/download).

In combination with the SARS-CoV-2 rRT-PCR (N-gene) targets, the CDC instructions list a table of expected results and associated interpretation that shows a decisive role of the human RNase P (h-RP) procedure control (Table 1).

**Table 1:** 2019-nCoV rRT-PCR Diagnostic Panel Results Interpretation Guide*.

| 2019 nCoV_N1 | 2019 nCoV_N2 | RP** | Result Interpretation | Report | Actions |
| --- | --- | --- | --- | --- | --- |
| + | + | ± | 2019-nCoV detected | Positive 2019-nCoV | Report results to CDC and sender. |
| - | - | + | 2019-nCoV not detected<br>↓<br>Invalid Result | Not Detected<br>↓<br>Invalid | Report results to sender. Consider testing for other respiratory viruses. |
| - | - | - | Invalid Result | Invalid | Repeat extraction and rRT-PCR. If the repeated result remains invalid, consider collecting a new specimen from the patient. |

\* Adapted from https://www.fda.gov/media/134922/download. Added in red are the interpretations that ought to be drawn based on the current CDC-recommended RP control: see main text.
\*\* Human RPP30 gene

During the set-up phase of this CDC-recommended SARS-CoV-2 rRT-PCR procedure in our lab, we noticed that the positive control for RNA extraction and/or RT efficacy, did not produce the expected results. According to the CDC instructions, the *Extraction Control* targets the human RNase P (RP) gene and a <40.00 Ct rRT-PCR value would indicate “…*the presence of the human RNase P gene.”*

In this study, we evaluate the CDC-recommended h-RP rRT-PCR control, identify its faulty design and offer an easy solution to correct it, without the need to completely change this positive control primer/probe set.

## RESULTS AND DISCUSSION

When we evaluated the h-RP-based rRT-PCR control, it became clear that the associated PCR primer/probe set is actually targeting the human MRP subunit p30 gene (h-RPP30, NM_001104546) of the Ribonuclease-P ribonucleoprotein complex.

Moreover, sequence analysis showed that both the CDC-RP forward rRT-PCR primer (CDC-RP-F), as well as the CDC-RP reverse rRT-PCR primer (CDC-RP-R) are completely located in exon 1 of the h-RPP30 gene (Figure 1). This disqualifies this primer set to be a proper positive control for a RT-PCR-based test, as without RNA and/or without proper RT synthesis, PCR amplification of even trace background genomic DNA (gDNA) would still result in a (false) positive signal^5^. Consequently, this faulty design has severe implications for the interpretation of the 2019-nCoV rRT-PCR results: in the case of absence of 2019 nCoV_N1/N2 PCR signals and presence of h-RP (h-RPP30) PCR signal, the conclusion “2019-nCoV not detected” should be “Invalid Result” (Table 1), as the PCR signal could originate from gDNA amplification.

**Figure 1:**
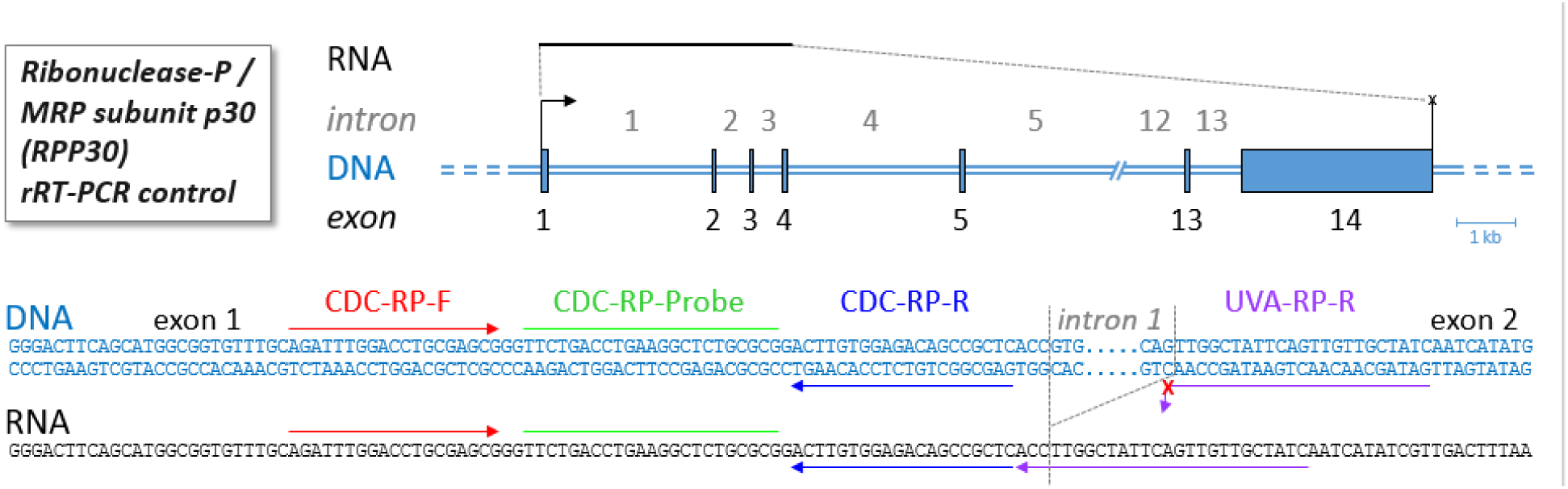
Positive-control rRT-PCR primer locations in DNA and RNA. Schematic representation of the exon/intron structure of the human RPP30 gene plus the locations of the CDC-recommended rRT-PCR primer/probe set (red: forward primer, green: probe, blue: reverse primer) and the alternative UvA-developed rRT-PCR primer (purple: reverse primer). Indicated is where the UvA-developed reverse primer does not match with the gDNA intron1-exon2 boundary sequence (red X).

To test whether our prediction that the CDC-recommended h-RPP30 rRT-PCR positive control can produce false-positive results, we performed a small experiment in which we compared the performance of the CDC-recommended h-RPP30 primer set on DNA and RNA. Strong rRT-PCR signals (= low Ct values) were obtained not only with RNA, but also with RNA free DNA as sample material (Table 2), which confirmed the ability of this primer set to amplify gDNA. To show the potential severity of this unwanted gDNA signal, we amplified four nose swab samples from two healthy individuals without and with RNase treatment to remove the RNA.

**Table 2:** Ct values for the signals from the human RP (h-RPP30) rRT-PCR analyses*.

| Sample | Treatment | CDC-recommended | UvA-adapted |
| --- | --- | --- | --- |
| Individual-1 left nostril |  | 23 | 31 |
| Individual-1 right nostril |  | 23 | 29 |
| Individual-2 left nostril |  | 25 | 25 |
| Individual-2 right nostril |  | 24 | 27 |
| Individual-1 left nostril | RNase | 23 | - |
| Individual-1 right nostril | RNase | 23 | - |
| Individual-2 left nostril | RNase | 27 | - |
| Individual-2 right nostril | RNase | 24 | - |
| Reference DNA (RNA free) sample 1 |  | 22 | - |
| Reference DNA (RNA free) sample 2 |  | 22 | - |
| Reference RNA (DNA free) sample 1 |  | 23 | 22 |
| Reference RNA (DNA free) sample 2 |  | 23 | 22 |
| Water sample 1 |  | - | - |
| Water sample 2 |  | - | - |

\* -, no rRT-PCR signal

One RNase-treated sample from Individual-2, showed for the CDC-recommended primer/probe set somewhat higher Ct values as the associated non-treated samples (Table 2), which reflects the contribution of h-RPP30 mRNA in the sample. In contrast, all other RNase-treated samples showed about identical Ct values as compared to the associated non-treated samples, which means that these signals by the CDC-recommended primer/probe set were entirely caused by amplification of gDNA. Other, similar experiments confirmed these findings (results not shown).

To prevent amplification from gDNA it is common practice^5^ in the design of rRT-PCR primer/probe sets to: i) design primer pairs in different exons, with large intron sequences in between, so the primers are too far apart in the gDNA, but close together in the spliced mRNA; or ii) design one or both primers/probe on exon-exon junctions, because these sequences are only present in spliced mRNA and not in gDNA. In order to repair the faulty CDC-recommended positive control rRT-PCR primer/probe set, yet change as few primers/probes as possible for practical reasons, we designed several new h-RPP30 rRT-PCR reverse primers according to the mentioned design principles, while keeping the CDC-recommended h-RPP30 (RP) forward primer and probe.

It turned out that the extra-intron-sequences approach in this case seemed less reliable, as we sometimes still got slight background signals with PCR-extension times of 60 seconds, even with PCR products well over 3,5 kb (results not-shown). Notwithstanding the limited design space, we were able to design an exon1-exon2 junction reverse primer (Figure 1) that performed good on RNA and not on gDNA. This University of Amsterdam (UvA)-developed reverse primer forms in combination with the CDC-recommended forward primer and probe, the UvA-adapted h-RPP30 rRT-PCR primer/probe set.

The UvA-adapted h-RPP30 primer/probe set was verified in a rRT-PCR experiment on the same samples as the CDC-recommended rRT-PCR set (Table 2). With the UvA-adapted rRT-PCR set, a somewhat better Ct value was obtained with the pure RNA samples and no signal with the pure gDNA samples. Thus, not only is the h-RPP30 UvA-adapted primer/probe set mRNA specific, it also slightly outperforms (∼1 Ct = 2-fold) the original CDC-recommended primer/probe set. These results were confirmed by the nose swab samples, of which all RNase-treated samples did not produce a rRT-PCR signal with this new primer/probe set. As the rRT-PCR signals from the UvA-developed primer/probe set of the nose swabs thus originate exclusively from mRNA amplification, they confirm our previous observation that there was a low amount of mRNA present in samples from Individual 1 (= high Ct values). The erroneous interpretation based on the CDC-recommended primer/probe, would be that the nose swab samples from Individual-1 would contain about three times (Ct difference = 1.5) more mRNA than those from Individual-2, whereas the UvA-adapted primer/probe set shows it to be actually an about 16 times reversed situation (Ct difference = −4).

## CONCLUDING REMARKS

We experimentally verified the faultiness of the h-RP positive control in the CDC-recommended 2019-nCoV rRT-PCR Diagnostic Panel caused by the design of the PCR primer pair in the same exon. It is somewhat puzzling that a primer/probe set that is frequently used to determine copy-number variation in the human genome, ended up as a control for RNA extraction and/or RT efficacy^6^, since it is clear from the design that background gDNA will pose a major problem.

We showed that the CDC-recommended primer/probe set indeed amplified gDNA and thus may lead to false interpretations and potentially false-negative 2019-nCoV diagnoses. Assuming a cut-off of Ct = 30, this would lead to a true (UvA-adapted) h-RPP30 absent measurement, yet a (strong) false (CDC-recommended) positive signal, due to background gDNA amplification, in one of our nose swab samples and thus a potentially wrong interpretation of the whole SARS-CoV-2 test. Admittedly, a cut-off of Ct = 30 is relatively low and in this particular nose swab sample just a very low amount of RNA might be present. Nevertheless, this sample illustrates the principle of false-positive results in that even without any extracted RNA there will be a strong signal in the positive control. Here the Ct difference between the mRNA amplification and gDNA amplification is already about 8 (= ∼256-fold).

A possible option to battle the faulty h-RPP30 primer/probe design, is to treat extracted sample RNA with DNase to degrade and eliminate background gDNA. However, in order to control for the total absence of gDNA, an extra RT-minus negative control then has to be added^4^, which seems not a very practical solution.

We therefore opted to just replace the reverse primer for this rRT-PCR positive-control and designed a exon-exon junction h-RRP30 reverse primer that in combination with the existing forward end probe would not amplify gDNA. By only replacing one primer of the faulty CDC h-RPP30 rRT-PCR primer/probe set, we were able to overhaul it into a mRNA specific primer/probe set with an at least equal amplification performance.

Given the worldwide importance of the widespread testing for the presence of the SARS-CoV-2 virus, we hope that the current CDC h-RP reverse primer will be replaced by all facilities that use the CDC-recommended SARS-CoV-2 rRT-PCR diagnostic panels as soon as possible, to further avoid potential false-negative COVID-19 diagnoses.

During the submission of our manuscript we discovered a study on the same topic by Adam P. Rosebrock with similar results^7^, but a different solution in that he proposes a completely new rRT-PCR h-RPP30 primer/probe set, whereas we only replace the reverse primer of the CDC-recommended primer/probe set.

## MATERIAL AND METHODS

### Biological materials and RNA isolation

Nasal swabs where taken from two adult healthy volunteers using STX 764 sterile small polyester swabs (Texwipe) and immediately stirred for 30 seconds in 1 ml of TRK lysis buffer (Omega Bio-tek). One volume of 70% EtOH was added, mixed by vortexing after which the mixture was loaded onto a spin column from the E.Z.N.A. Total RNA Kit I (Omega Bio-tek). The column was washed according the manufacturer’s instructions and the purified nucleic acids were eluted in 40 μl nuclease-free water. Half of the eluate (20 μl) was treated with 1 μl RNase Cocktail Enzyme Mix (Thermo Fisher Scientific) at 37°C for 30 min.

Universal Human Reference RNA (Stratagene) was treated with RNase-Free DNase Set (Qiagen) according to the manufacturer’s protocol to remove possible traces of genomic DNA.

### Real-time reverse-transcription PCR (rRT-PCR)

The 2019-nCoV CDC RUO Primers and Probes (IDT) were used for the CDC-recommended h-RPP30 assay. The UvA-modified primer and probe set makes use of the original CDC forward primer (CDC-RP-F) and probe (CDC-RP-probe). The reverse primer was redesigned by the UvA (UVA-RP-R) and ordered at IDT: gatagcaacaactgaatagccaaggt.

cDNA synthesis and real-time PCR were combined in a one-step PCR reaction (rRT-PCR) using the TaqPath 1-Step RT-qPCR Master Mix, CG (Thermo Fisher Scientific) on a QuantStudio 3 Real-Time PCR System (Thermo Fisher Scientific).

rRT-PCR reaction set-up and thermal cycling conditions were taken from the original CDC-reccomended protocol (https://www.fda.gov/media/134922/download).

Nucleic acid templates were either 100 ng DNAse-treated Universal Human Reference RNA, 500 ng human genomic DNA (Promega) or 5 μl of untreated or RNase-treated nasal swab nucleic acid isolates. 5 μl of nuclease-free water was used as no-template controls.

## CONFLICT OF INTEREST STATEMENT

The Authors declares that there is no conflict of interest.

## Notes

### Competing Interest Statement

The authors have declared no competing interest.

### Summary of Updates

Relevant study discovered during submission, small paragraph concerning this study was added in the concluding remarks and the related reference added to the references.

